# Integrated Biomedical System

**DOI:** 10.1101/050138

**Authors:** Darrell O. Ricke, James Harper, Anna Shcherbina, Nelson Chiu

## Abstract

Capabilities for generating and storing large amounts of data relevant to individual health and performance are rapidly evolving and have the potential to accelerate progress toward quantitative and individualized understanding of many important issues in health and medicine. Recent advances in clinical and laboratory technologies provide increasingly complete and dynamic characterization of individual genomes, gene expression levels for genes, relative abundance of thousands of proteins, population levels for thousands of microbial species, quantitative imaging data, and more - all on the same individual. Personal and wearable electronic devices are increasingly enabling these same individuals to routinely and continuously capture vast amounts of quantitative data including activity, sleep, nutrition, environmental exposures, physiological signals, speech, and neurocognitive performance metrics at unprecedented temporal resolution and scales. While some of the companies offering these measurement technologies have begun to offer systems for integrating and displaying correlated individual data, these are either closed/proprietary platforms that provide limited access to sensor data or have limited scope that focus primarily on one data domain (e.g. steps/calories/activity, genetic data, etc.). The Integrated Biomedical System is being developed to demonstrate an adaptable open-source tool for reducing the burden associated with integrating heterogeneous genome, interactome, and exposome data from a constantly evolving landscape of biomedical data generating technologies. The Integrated Biomedical System provides a scalable and modular framework that can be extended to include support for numerous types of analyses and applications at scales ranging from personal users, communities and groups, to large populations.

**Disclaimer:** This work is sponsored by the Assistant Secretary of Defense for Research & Engineering under Air Force Contract #FA8721-05-C-0002. Opinions, interpretations, recommendations and conclusions are those of the author and are not necessarily endorsed by the United States Government.

## Introduction

Human health and performance is understood to be affected by both nature (genome) and nurture (activities & environment). One notable example of the combined effects of genetics and the environment on health is the identification that the GRIN2A gene significantly modulates risk for developing Parkinson’s disease, but only in heavy coffee-drinkers [1]. This study provides proof that inclusion of quantitative measures of environmental factors can help identify important genes that would be otherwise missed in GWAS studies that ignore exposures. However, the challenges associated with designing and implementing broad quantitative studies of complex interactions at scales sufficient to achieve sufficient statistical power are considerable.

There are multiple efforts underway that are making progress toward addressing the challenges of integrating genome, interactome, and exposome[2] data to support focused scientific studies. The Institute of Systems Biology’s Hundred Person Wellness Project[3] and 100K Project[4] are integrating genomics, monitors, and blood sampling to build on the pioneering N-of-one work conducted by Larry Smarr[5] and Michael Snyder[6, 7] to articulate the vision and promise of predictive, preventative, personalized, and participatory (P4) medicine[8]. Orion Bionetworks[9] is combining traits, genetics, and interactome with a focus on brain disorders. Sanchez *et al*. [10] has also proposed exposome informatics integrating the genome, phenome, and exposome. Systems integrating personal sensors and exposome have been developed by Doherty & Oh[11] and Nieuwenhuijsen *et al*. [12]. Other relevant available resources include PhysioNet[13] and MOPED[14]. The Human Longevity project[15] is examining genome, microbiomes, and metabolites of volunteers. While these projects all share the common elements of longitudinal integration of heterogeneous biomedically relevant data, each either focuses on a relatively narrow set of measurements or relies on custom data storage and analysis architectures that do not provide a scalable foundation for larger-scale integration across studies to enable meta-analysis of data from multiple studies.

The Integrated Biomedical System is being developed as an open source platform for integrating genome, interactome, and exposome data (see Figures 1 & 2) that provides a unifying model to promote more open data sharing and analysis. The software architecture with multi-scale operability design intended to scale from running on a single laptop/workstation as a standalone system with an embedded private local database, to a study platform, to large-scale implementations all using standard scalable web technology stacks.

**Figure 1.**
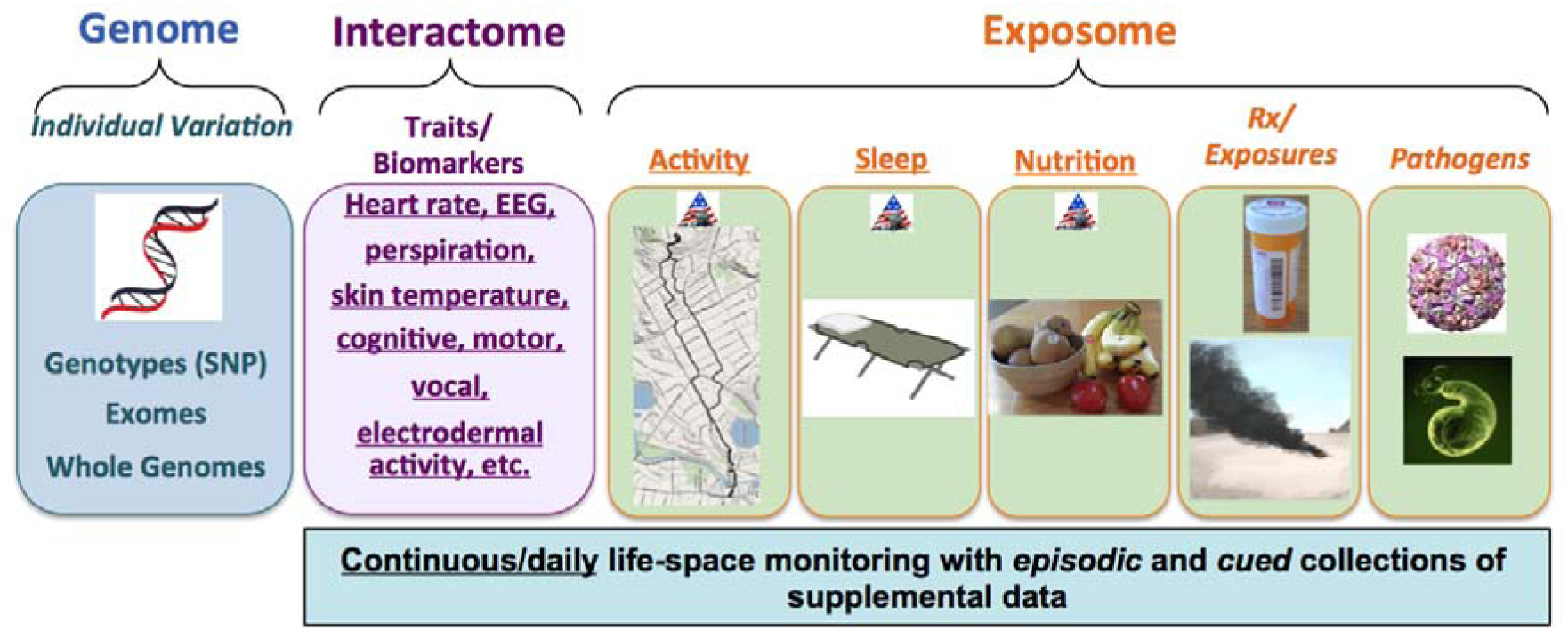
Integrated Biomedical System Vision. Integrating genome, interactome, and exposome heterogeneous data to create an open data system to promote health, wellness, and future biomedical discoveries.

**Figure 2.**
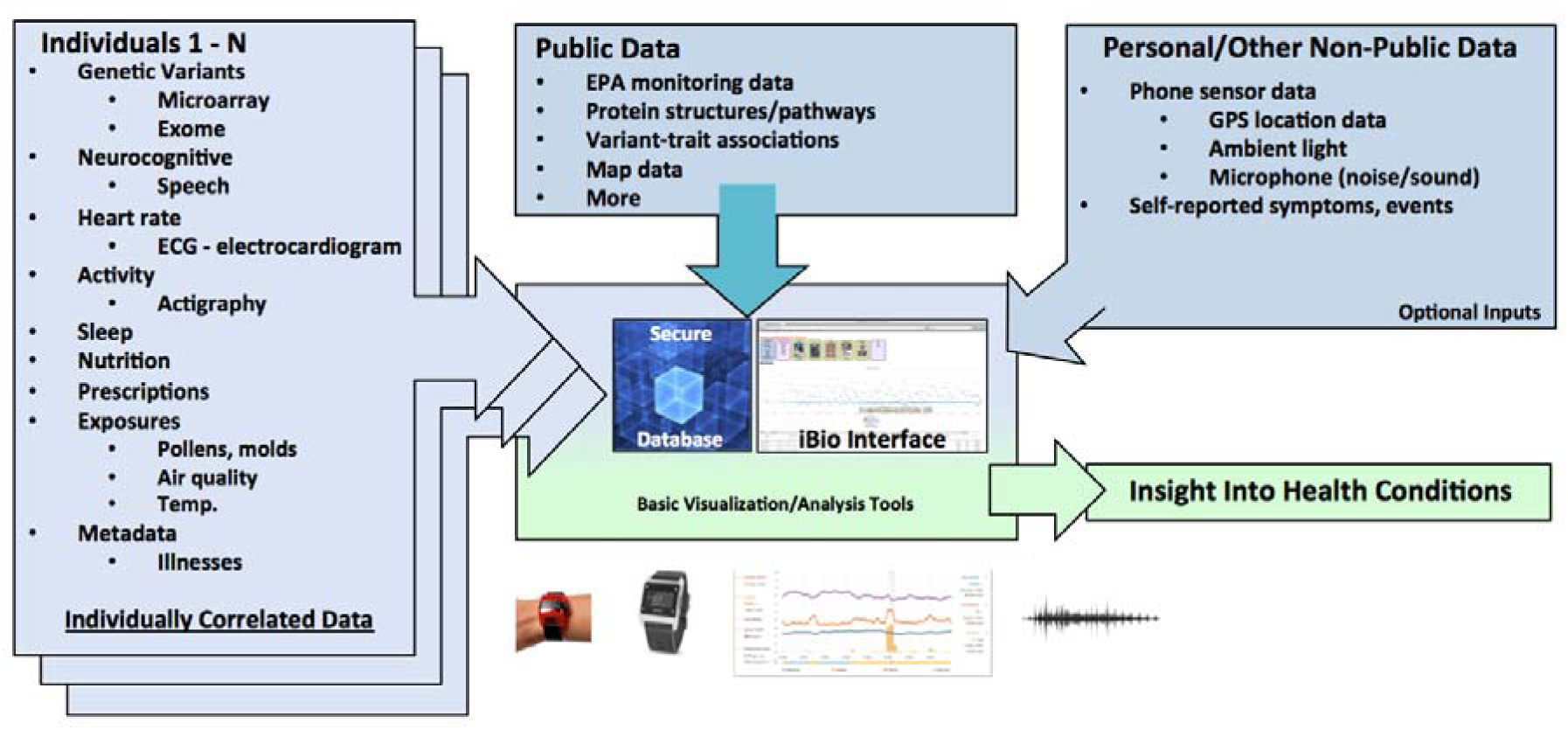
Data Integration and Analysis. Our vision of organizing data from individuals, electronic health records, and public data sources into a platform for integrating heterogeneous data sources for data analysis and correlation mining.

## Methods

### Protocol design & approvals

The Integrated Biomedical System description and written consent form was reviewed and approved by the MIT Committee on the Use of Humans as Experimental Subjects (COUHES) for volunteers.

### Integrated Biomedical System (iBio)

The Integrated Biomedical System was developed on the Ruby on Rails[16] platform with Ruby gems and JavaScript plugins. The Rails platform supports multiple SQL relational databases including MySQL and no SQL databases such as Mongo DB. MySQL, Oracle, Mongo DB, etc. all scale to over a billion records in a single table. The underlying architecture and approach can be extended to handle a variety of additional data sources. The Integrated Biomedical System and Rails can be installed on computers ranging from stand-alone on a laptop/desktop to servers running Windows OS, Mac OS, Linux, or Unix. Individuals can install and run this system for personal use without needing to set up a web service; to facilitate this the default configuration uses the Sqlite3 database, which installs with the Rails setup. Switching to MySQL or Oracle requires database software installation and a 5 line update to the Rails database.yml configuration file with updated database instance details. To facilitate bulk loading of large numbers of data files, command line interfaces for each ETL module are included in the app/utilities folder.

Extract, transform, and load (ETL) modules were developed for 23andMe SNPs[17] files, SwissProt[18] dat file, DrugBank[19] XML file, NCBI Gene[20] gene file, PharmGKB pathways[21], and Protein Data Bank (PDB) protein structures [22]. After SwissProt sequences and PDB protein structures were loaded, the structure coordinates were mapped to sequence residues with the included app/utilities/align_pdb.rb tool; this enables the visualization of residues and variants on structures. Interface modules were developed to allow individual or pooled variants to be visualized on protein structures with the integrated Jmol[23] structure viewer.

### Interactome

Interactome data included in the pilot collection described herein includes heart rate, interbeat interval (IBI), and electrocardiogram (ECG), skin temperature, skin conductance, galvanic skin response, and respiratory rate. These aggregated data were collected by a diverse collection of commercially available wearable physiological monitoring devices. All volunteers were offered a Basis B1 watch[24] and Polar Loop H7 heart rate monitor[25]. A subset of volunteers are evaluating Hildago Equivital EQ-02-SEM[26], Empatica E3[27], Mio Link[28], and Zephyr BioHarness 3[29] devices. Data logging functionality was not built in for the Polar Loop and Mio Link heart rate monitors, so these data streams were wirelessly synced and stored continuously on co-worn Actigraph Actisleep device. ETL modules were developed for Basis B1 json files[30], Actigraph heart rate csv or dat files (including Polar Loop and Mio Link), Empatica E3 zip files, Hidalgo Equivital SEM2 persisted summary csv files, Zephyr BioHarness summary csv files, vocal recordings and associated Matlab.mat files. Data displays include Ruby gems and JavaScript plugins: Google Maps[31], jQuery[32], lazy_high_charts[33], Highstocks[34], Data-Drive Documents (D3)[35], FullCalendar[36], rails3-jquery-autocomplete[37], and more. The graphical user interface for “ Data Loading” provides the ability to download data from the Basis web site and drag and drop interfaces for easy file uploads for each of the device ETL modules.

### Exposome

Activity and sleep were monitored continuously using wearable and personal electronic devices that used algorithms to process raw data provided by built-in 3-axis accelerometers. Data describing daily nutrition, prescriptions, and over-the-counter medications were collected manually and provided by a subset of volunteers. Devices used by volunteers for continuous data collection included the Fitbit Flex, the Basis B1 watch, Actigraph ActiSleep monitors, basic Actigraph activity monitors GT3X+, Jawbone Up, and smart-phone apps including MyTracks, and Sleep Cycle. ETL modules were developed for Fitbit csv files, Jawbone csv files, Actigraph[38] sleep csv files, MyTracks app[39] csv files, and Sleep Cycle app[40] csv files. Additionally to demonstrate the ability to integrate other publicly available data, modules were developed for integration of EPA AirData (daily and hourly csv files[41]), and foods[42]. Graphical user interfaces were developed for entering activities, events, meals, drinks, prescriptions, and over-the-counter medicines. Multiple volunteers submitted oral swab samples for metagenomics sequence analysis when sick (cued data collection).

## Results

### Interactome

#### Heart Rate Monitoring

Heart rate monitoring devices provide heart rate, interbeat interval (IBI), and electrocardiogram (ECG) measurements. Heart rate measurements for multiple devices for an individual are shown in Figure 3. Hidalgo Equivital SEM2 and Zephyr BioHarness were typically worn only during more active periods. Lower Zephyr heart rate values observed on Aug. 29 likely resulted from the contact pads drying out during a period of extended wearing with low activity level. Some data gaps result from the need for device battery recharging (Empatica E3 - daily and Mio Link every 8 to 10 hours). Higher correlations of results are observed for periods of sleeping and light activity. This observation is consistent with previous anecdotal observations of data accuracy and coverage decreases for many wearable sensors during periods of high activity.

**Figure 3.**
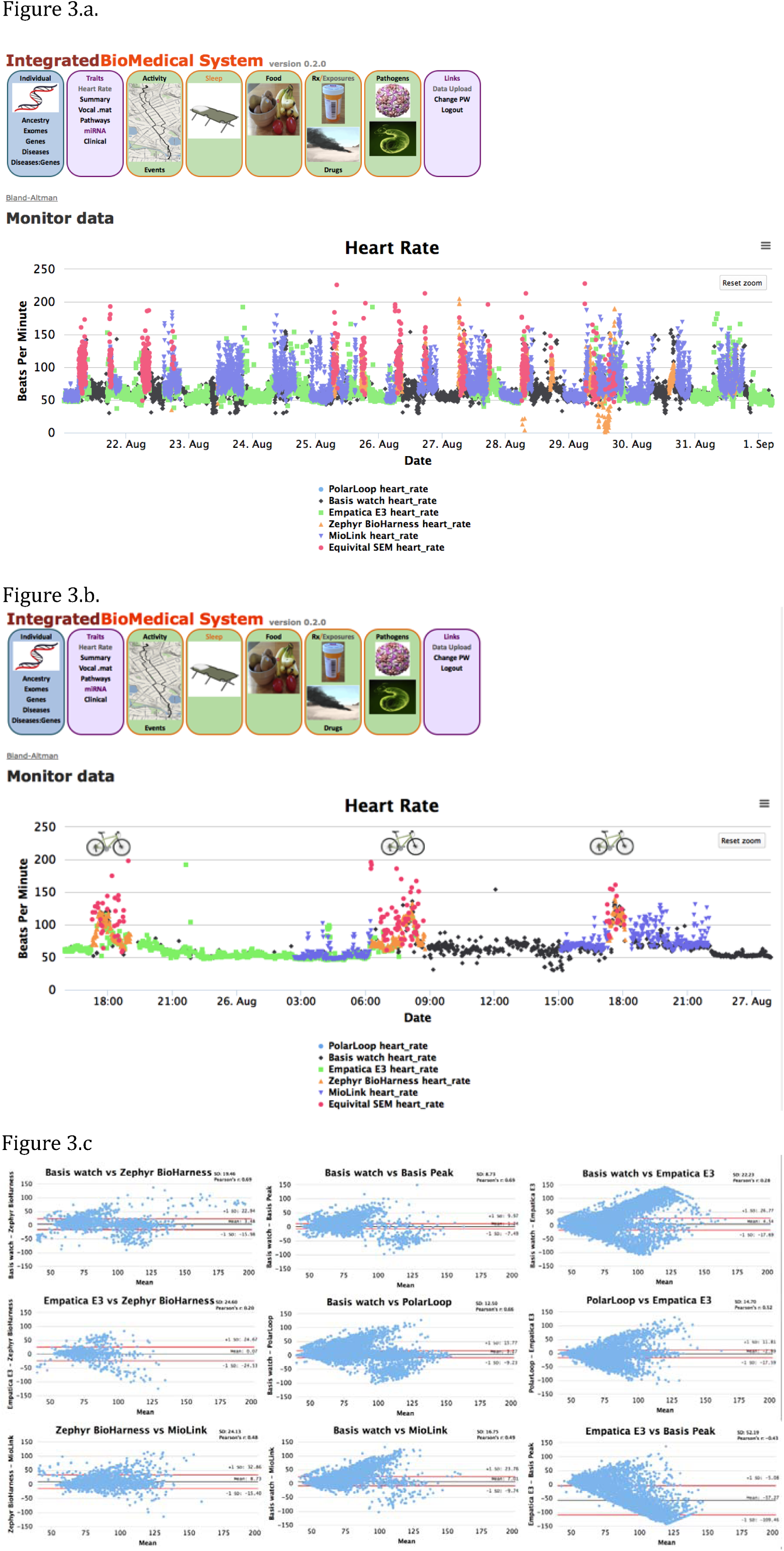
Heart Rate Monitoring. (a) Screen shot of heart rate beats per minute measurements for a volunteer wearing Basis B1 watch, Empatica E3, Zephyr BioHarness, Hildago Equivital SEM2, and Mio Link devices. SEM2 values were filtered for minimum quality values of 70 with selection of median value. (b) Zoomed in view of heart rates illustrating measurements at different activity levels. (c) Bland-Altman plots comparing measurements from the heart rate tracking devices with corresponding Pearson r correlation values.

### Exposome

#### Sleep Monitoring

Multiple devices tested provide top-level estimates of nightly time asleep and number of sleep interruptions. Some devices also attempt to break down the sleep time into sleep phases (light, deep, and rapid eye movement - REM sleep). This data was integrated to enable comparisons of sleep classifications assigned by these devices (investigation of the accuracy of these estimates vs. gold-standard polysomnography was beyond the scope of the present work). Example longitudinal measurements from a single individual collecting data in parallel using Jawbone Up, Basis B1, Fitbit, and ActiSleep are shown in Figure 4. Analytical modules enabling pairwise comparisons of unfiltered nightly time asleep estimates between different devices were developed and integrated into the Integrated Biomedical System. Simple comparisons of daily total time asleep reported across the range of devices revealed a lack of correlation for most device pairs as measured by Pearson r statistics. Likewise, finer-grained estimates of light sleep (provided by Basis and Jawbone) and deep sleep (Jawbone) compared to deep sleep plus REM sleep (Basis) were also poorly correlated. Only the two Actigraph algorithms, Sadeh and Cole-Kripke, which were run on the same raw Actigraph sensor data produced highly correlated results (r of 0.97).

**Figure 4.**
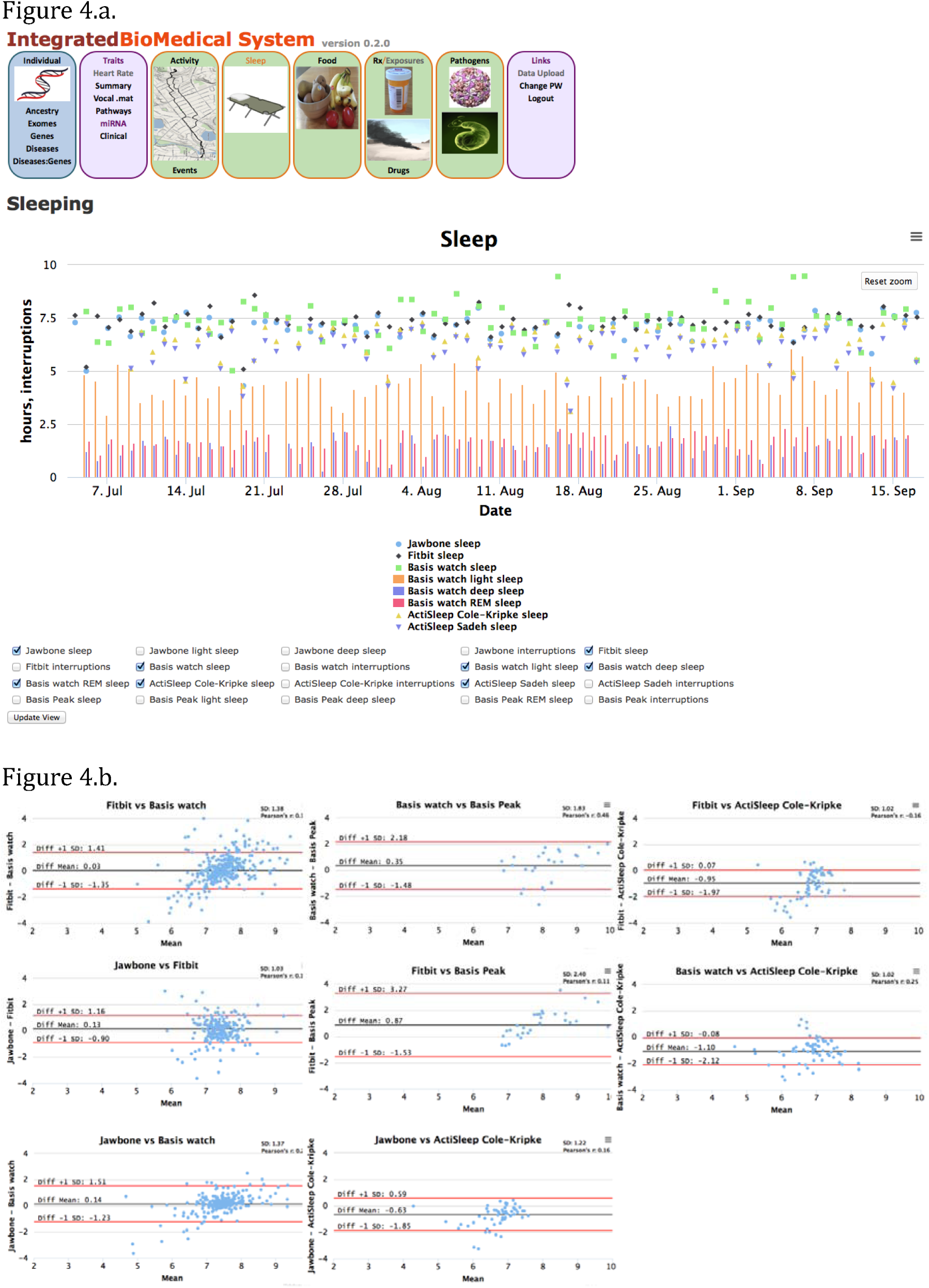
Sleep Monitoring. (a) Screen shot of daily total sleep measurements for a volunteer for Fitbit Flex, Jawbone Up, Basis B1 watch, and Actisleep. (b) Bland-Altman plots comparing measurements from the sleep tracking devices for this volunteer.

### Exposures

Global Position System (GPS) tracking of outside activities available in the Integrated Biomedical System from smartphone or GPS data can provide continuous localization for an individual. This data enables a range of potentially useful correlations to be determined including correlations with data from nearby EPA or other air quality monitoring station(s) as an initial step toward quantitative tracking of individual exposures. Inferred exposure levels can be estimated from nearby sensors for a wide variety of measured pollutants, particulates[41], and pollen levels[43]. Figure 5 illustrates NO2, PM2.5, and PM10 exposures for an afternoon walk.

**Figure 5.**
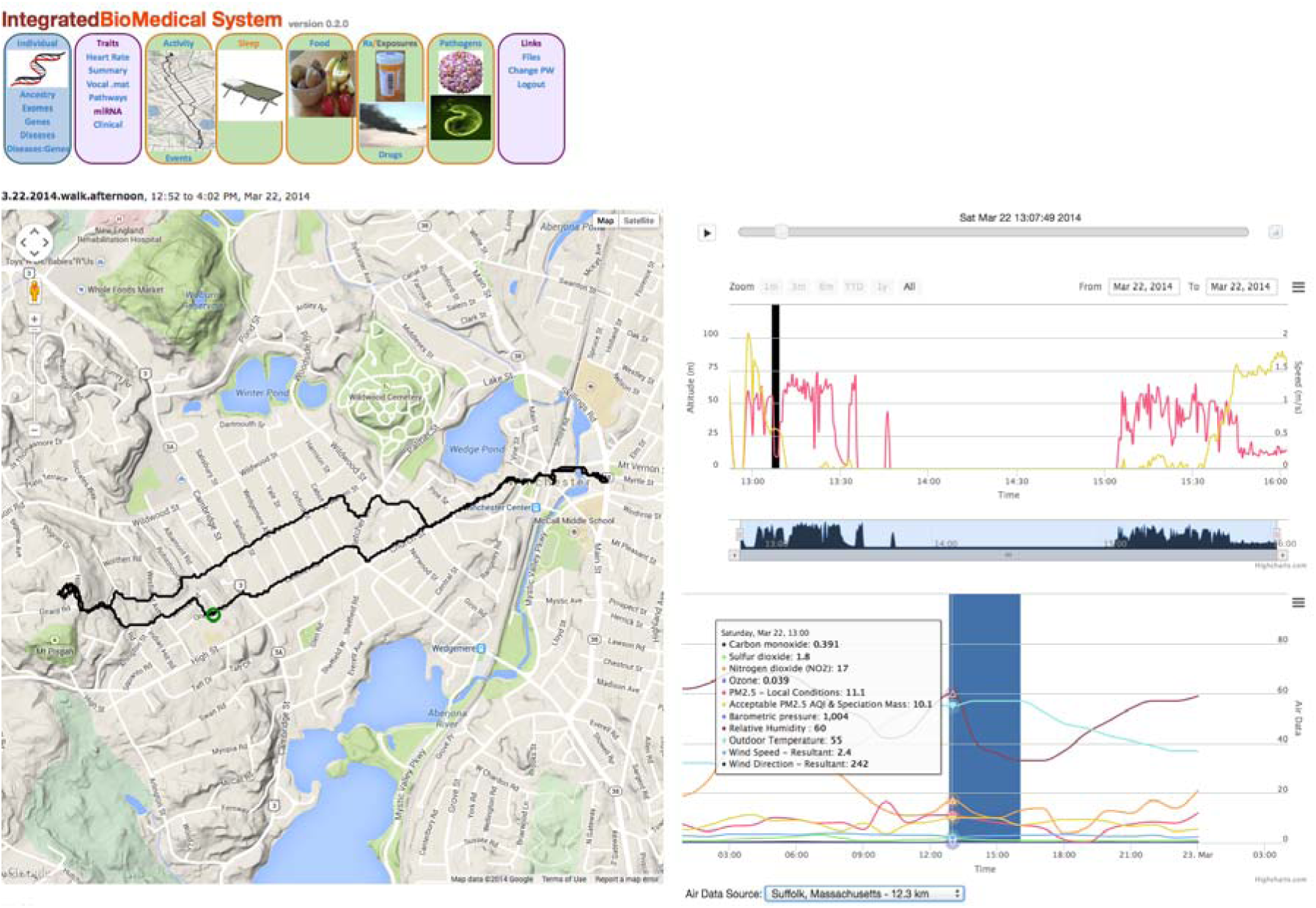
Outdoor walk and Integration with EPA AirData. Example visualization of activity data with estimated exposure levels from nearby EPA AirData monitoring site.

## Discussion

### Vision

Genome, interactome, and exposome all influence an individual’s wellness. The Integrated Biomedical System was developed to demonstrate the ability to begin integrating these heterogeneous data sources in near real-time for individuals. This was accomplished using an architecture that can operate on a stand-alone laptop or desktop personal computer (PC) to provide additional privacy and security and can be connected seamlessly to voluntarily transfer selected data to centralized highly scalable systems built on the same data architecture that can integrate data from many thousands or even millions of individuals. This approach could provide a path to developing new crowd-sourced models for large-scale prospective/retrospective studies of how individual combinations of genomic and environmental factors correlate with a range of human health and performance traits. Individual monitoring devices, genetic data, blood biochemistries, nutrition, exposures, illnesses, vocal and additional data have been organized and integrated into a unified system (see Figure 2). Using the same tools and architectures, additional quantitative lab results and diagnostic data like images and physiological monitoring system data can be added to further increase the research scope of the system. Incorporation of additional natural language processing tools and data architecture modifications can enable text-based metadata collections (e.g. regular symptoms logging from personal health blogs, social interaction details from social media platforms, information from electronic health records) to be included in future versions of the system. Furthermore, these personal datasets can be combined with relevant public datasets and other non-public data to provide new insights into health-associated effects to support detailed N-of-1 and population retrospective analyses.

### Genome

As large scale DNA sequencing costs continue to decrease, sequencing an individual’s DNA becomes more affordable and practical. While tools exist to characterize variants (Polyphen2, SIFT, etc.), the potential to correlate variants with protein structure/function, physiology, molecular biomarkers, etc. typically is done manually and within studies with a single focus. Integrating genomic data with interactome and exposome data will help create new opportunities for turning data into new discoveries and knowledge. As advances in DNA sequencing technology enable more widespread access to genomic data for individuals, the ability to correlate that data with quantitative interactome and exposome data will become increasingly important. Together, these data can broadly enable efforts to elucidate the interplay between genomic and environmental factors that contribute to complex individual human traits and health.

### Interactome

Cognitive performance and health phenotypes can be assessed through a variety of indirect methods including analysis of biomarkers in blood, psychomotor vigilance task (PVT), profile of mood states (POMS), automated neuropsychological assessment metrics (ANAM), speech analysis, facial and eye movement tracking, electroencephalography (EEG), and similar approaches. These assessments and others have been developed and used quantitatively define progressions of important traits/symptoms in individuals experiencing a number of conditions including depression[44], posttraumatic stress disorder (PTSD), and traumatic brain injury (TBI), as well as environmental stressors including sleep disruption, etc. Data streams produced from these assessments combined with traditional measurements of traits, molecular biomarkers, and clinical data to provide a new platform for gaining insight into the underlying physiology individual health, fitness, and well being. Retrospective analysis of large-scale collections will provide future biomedical discoveries. Increasing proportions of future biomedical discoveries will be driven by the ability to effectively collect, manage, and interpret massive amounts of heterogeneous data. Enhancements to integrate additional interactome data types and analysis tools are currently underway and these features will be included in future releases.

### Exposome

Asthma and COPD affect 18.7 and 6.8 million individuals in the United States[45]. Environmental exposures can exacerbate these conditions[46]. Asthma can be triggered by particulate matter, ozone, sulfur dioxide, nitrogen oxide, and pollens[47]. Devices, including smart phones, with GPS tracking ability enable the possibility of data integration with environmental monitoring data. Nearby monitoring stations and mobile monitoring devices provide weather and exposure estimates that can be correlated using time stamped GPS positional information. Monitoring stations track a rich variety of environmental exposure data[41]. While the current system provides incomplete coverage, it demonstrates a viable path to incorporation of additional sensor streams (including indoor air quality sensors, UV exposures, etc.) and activity-based estimates of indoor vs. outdoor exposures. It will be possible to provide increasingly complete individualized and integrated quantitative estimates of specific exposures that can be correlated with possible health effects, symptoms, and well-being. Larger and more complete data sets enabled by integrated systems like the one described here, can play a key enabling role for more quantitative genome vs. environment studies in the future.

### Looking Forward

Health and fitness are impacted by genetics, interactome, and exposome. Solutions that combine relevant data across these data domains will lead to new health and fitness insights. The data infrastructure to collect and aggregate data across these domains is currently lacking but large corporations are moving rapidly in this direction with cloud-based private solutions. These corporate solutions provide access to summary data (steps walked, hours slept, etc.), but rarely access to the underlying raw data. Typically, these systems require users to consent to granting data ownership to the corporation and not themselves. Open data architectures with open source solutions can provide alternatives to individuals and organizations for personal health, fitness, and wellness promotion and also longitudinal studies to facilitate data exploitation for research and discoveries and decision support for leaders and medical personnel. The Integrated Biomedical System is available on GitHub [https://github.com/doricke/IBio].

## Acknowledgements

The authors would like to acknowledge Carl Ricke for graphic artwork.

## References

1. Hamza, T.H., et al., Genome-Wide Gene-Environment Study Identifies Glutamate Receptor Gene GRIN2A as a Parkinson’s Disease Modifier Gene via Interaction with Coffee. PLOS Genetics, 2011. 7(8).

2. Wild, C.P., Complementing the Genome with an “ Exposome”: The Outstanding Challenge of Environmental Exposure Measurement in Molecular Epidemiology. Cancer Epidemiology Biomarkers & Prevention, 2005. 14(8): p. 1847–1850.

3. Gibbs, W.W., Medicine gets up close and personal. Nature, 2014. 506(7487): p. 114–115.

4. Hood, L. and N.D. Price, Demystifying Disease, Democratizing Health Care. Sci Transl Med, 2014. 26.

5. Smarr, L., Quantifying your body: A how-to guide from a systems biology perspective. Biotechnology Journal, 2012. 7(8): p. 980–991.

6. Li-Pook-Than, J. and M. Snyder, iPOP goes the world: integrated Personalized Omics Profiling and the road towards improved health care. Cell Biol., 2013. 20(5): p. 660–666.

7. Chen, R. and M. Snyder, Systems biology: personalized medicine for the future? Current Opinion in Pharmacology, 2012. 12(5): p. 623–628.

8. Hood, L. and C. Auffray, Participatory medicine: a driving force for revolutionizing healthcare. Genome Medicine, 2013. 5.

9. Xu, X., et al., Structural Characterization of the 1918 Influenza Virus H1N1 Neuraminidase. Journal of Virology, 2008. 82(21): p. 10493–10501.

10. Sanchez, F.M., et al., Exposome informatics: considerations for the design of future biomedical research information systems. J. Am Med Inform Assoc, 2014. 21(3): p. 386–390.

11. Doherty, S.T. and P. Oh, A Multi-Sensor Monitoring System of Human Physiology and Daily Activities. Telemedicine and e-Health, 2012. 18(3): p. 185–192.

12. Nieuwenhuijsen, M.J., et al., Using Personal Sensors to Assess the Exposome and Acute Health Effects. Int. J. Environ. Res. Public Health, 2014. 11: p. 7805–7819.

13. Goldberger, A.L., et al., PhysioBank, PhysioToolkit, and PhysioNet: Components of a New Research Resource for Complex Physiologic Signals. Circulation, 2000. 101(23): p. e215–e220.

14. Montague, E., et al., MOPED 2.5—an integrated multi-omics resource: multi-omics profiling expression database now includes transcriptomics data. OMICS, 2014. 18(6): p. 335–343.

15. Darwin, C., On the Origin of Species. 1859.

16. Samani, N.J., M. Tomaszewski, and H. Schunkert, The personal genome-the future of personalised medicine? The Lancet. 375(9725): p. 1497–1498.

17. Ormond, K.E., et al., Challenges in the clinical application of whole-genome sequencing. The Lancet. 375(9727): p. 1749–1751.

18. Boeckmann, B., et al., The SWISS-PROT protein knowledgebase and its supplement TrEMBL in 2003. Nucleic Acids Research, 2003. 31(1): p. 365–370.

19. Law, V., et al., DrugBank 4.0: shedding new light on drug metabolism. Nucleic Acids Research, 2014. 42(D1): p. D1091–D1097.

20. Just, W., Computational Complexity of Multiple Sequence Alignment with SP-Score. Journal of Computational Biology, 2001. 8(6): p. 615–623.

21. Wang, L. and T. Jiang, On the complexity of multiple sequence alignment. J. Comput. Biol., 1994. 1(4): p. 337–348.

22. Berman, H.M., et al., The Protein Data Bank. Nucleic Acids Research, 2000. 28(1): p. 235–242.

23. Prosite database. Available from: http://prosite.expasy.org/.

24. Fitch, W.M. and C.H. Langley, Protein evolution and the molecular clock. Fed. Proc., 1976. 35(10): p. 2092–2097.

25. Polar Loop H7 heart rate sensor. Available from: http://www.polar.com/us-en/products/accessories/H7_heart_rate_sensor.

26. Ricke, D.O. BioTools. Bioinformatics programs]. Available from: https://github.com/doricke/BioTools.

27. Gribskov, M., A.D. McLachlan, and D. Eisenberg, Profile analysis: detection of distantly related proteins. Proceedings of the National Academy of Sciences of the United States of America, 1987. 84(13): p. 4355–4358.

28. Mio Link heart rate band. Available from: http://www.mioglobal.com/Default.aspx.

29. Jmol: an open-source Java viewer for chemical structures in 3D. Available from: http://www.jmol.org.

30. Sigrist, C.J.A., et al., PROSITE: A documented database using patterns and profiles as motif descriptors. Briefings in Bioinformatics, 2002. 3(3): p. 265–274.

31. Whittle, J.R.R., et al., Broadly neutralizing human antibody that recognizes the receptor-binding pocket of influenza virus hemagglutinin. Proceedings of the National Academy of Sciences, 2011. 108(34): p. 14216–14221.

32. Consortium, T.U., Activities at the Universal Protein Resource (UniProt). Nucleic Acids Research, 2014. 42(D1): p. D191–D198.

33. Ricke, D.O., Analysis of Sequence and Molecular Evolution Information in Two Model Systems. 1995, Mayo Graduate School.

34. Bottema, C.K., et al., The pattern of spontaneous germ-line mutation: relative rates of mutation at or near CpG dinucleotides in the factor IX gene. Human Genetics, 1993. 91(5): p. 496–503.

35. Koeberl, D.D., et al., Functionally important regions of the factor IX gene have a low rate of polymorphism and a high rate of mutation in the dinucleotide CpG. Am J Hum Genet, 1989. 45(3): p. 448–457.

36. Povolotskaya, I.S. and F.A. Kondrashov, Sequence space and the ongoing expansion of the protein universe. Nature, 2010. 465(7300): p. 922–926.

37. Ashley, E.A., The precision medicine initiative: A new national effort. JAMA, 2015.

38. Ricke, D.O., Divergence Model of Protein Evolution. submitted.

39. Google MyTracks app. Available from: https://play.google.com/store/apps/details?id=com.google.android.maps.mytracks&hl=en.

40. Sleep Cycle alarm clock app. Available from: http://www.sleepcycle.com/.

41. Sommer, S.S. and R.P. Ketterling, The factor IX gene as a model for analysis of human germline mutations: an update. Human Molecular Genetics, 1996. 5(Supplement 1): p. 1505–1514.

42. The Hacker’s Diet. Available from: https://http://www.fourmilab.ch/.

43. Gribskov, M., A.D. McLachlan, and D. Eisenberg, Profile analysis: detection of distantly related proteins. Proceedings of the National Academy of Sciences, 1987. 84(13): p. 4355–4358.

44. Williamson, J.R., et al., Vocal biomarkers of depression based on motor incoordination, in Proceedings of the 3rd ACM international workshop on Audio/visual emotion challenge. 2013, ACM: Barcelona, Spain. p. 41–48.

45. Sahini, L., A. Tempczyk-Russell, and R. Agarwal, Large-Scale Sequence Analysis of Hemagglutinin of Influenza A Virus Identifies Conserved Regions Suitable for Targeting an Anti-Viral Response. PLoS ONE, 2010. 5(2): p. e9268.

46. Ko, F.W.S. and D.S.C. Hui, Air pollution and chronic obstructive pulmonary disease. Respirology, 2012. 17(3): p. 395–401.

47. Guarnieri, M. and J.R. Balmes, Outdoor air pollution and asthma. The Lancet. 383(9928): p. 1581–1592.

